# Differential Expression of Mucins in Murine Olfactory Versus Respiratory Epithelium

**DOI:** 10.1101/620757

**Authors:** Christopher Kennel, Elizabeth A. Gould, Eric D. Larson, Ernesto Salcedo, Thad W. Vickery, Diego Restrepo, Vijay R. Ramakrishnan

## Abstract

Mucins are a key component of the airway surface liquid and serve many functions. Given the numerous differences in olfactory versus respiratory nasal epithelia, we hypothesized that mucins would be differentially expressed between these two areas. Secondarily, we evaluated for changes in mucin expression with radiation exposure, given the clinical observations of nasal dryness, altered mucus rheology, and smell loss in radiated patients. Immunofluorescence staining was performed in a mouse model to determine the expression of mucins 1, 2, 5AC and 5B in nasal respiratory and olfactory epithelia of control mice and one week after exposure to 8 gy of radiation. Mucins 1, 5AC and 5B exhibited differential expression between olfactory and respiratory epithelium while mucin 2 showed no difference. Within the olfactory epithelium, mucin 1 was located in a lattice-like pattern around gaps corresponding to dendritic knobs of olfactory sensory neurons, whereas in respiratory epithelium it was only intermittently expressed. Mucin 5AC was expressed by subepithelial glands in both epithelial types but to a higher degree in the olfactory epithelium. Mucin 5B was expressed by submucosal glands in the olfactory epithelium but by surface epithelial cells in respiratory epithelium. At one-week after exposure to single-dose 8 gy of radiation, no qualitative effects were seen on mucin expression. Our findings demonstrate that murine olfactory and respiratory epithelia express mucins differently, and characteristic patterns of mucins 1, 5AC, and 5B can be used to define the underlying epithelium. Radiation (8 gy) does not appear to affect mucin expression at one week.

**Author Roles:** Christopher Kennel conceived, organized and executed the study, performed the analysis, and contributed to the manuscript.

Elizabeth Gould conceived and executed the study, and contributed to the manuscript.

Diego Restrepo conceived and executed the study, supervised the experiments, reviewed the analysis, and contributed to the manuscript.

Ernesto Salcedo performed experiments and reviewed the manuscript.

Thad Vickery performed experiments and reviewed the manuscript.

Eric Larson performed experiments and reviewed the manuscript.

Vijay Ramakrishnan conceived and executed the study, reviewed the analysis, and contributed to the manuscript.

All authors discussed the results and implications and contributed to the final manuscript.

## Introduction

Mucins are large glycoproteins (up to 1,500 nm in length) found in the hydrophilic gel of mucus overlying the airway epithelium (Rose and Voynow 2006; Hattrup and Gendler 2008). Mucus serves as a protective barrier against desiccation and a wide range of foreign substances including chemical irritants, particulates, and bacterial, fungal and viral pathogens. Both the remarkable size and extensive glycosylation of mucins contribute to their function in the airways where they protect from the continuous exposure to airborne bacterial and viral pathogens, toxins, and contaminants. *In vivo*, mucus contains over 90% water, and mucins constitute up to 80% of the dry weight making them the major protein component of mucus (Lai et al. 2009).

Mucins are a key component of the mucus overlying the sinonasal epithelium. Of the 21 known mucin isoforms, mucins 1, 4, 5AC and 8 have been found in human sinonasal epithelium and mucins 1, 5B, and 8 in sinonasal glands (Ali and Pearson 2007; Martinez-Anton et al. 2006; Rubin 2010; Cone 2005; Cone 2009). Importantly, some mucins, including mucin 1, have a transmembrane spanning peptide that binds them to cell membrane while other mucins are entirely secreted (Hattrup and Gendler 2008). The point of contact between the shorter and lower viscosity, membrane-bound mucins and the higher viscosity secreted mucins forms a slippage plane and double-barrier against environmental insults (Cone 2005; Cone 2009). Under stress, the secreted mucus layer may entirely shear away from the surface-bound mucus layer, which remains bound to the underlying cells (Lai et al. 2009; Cone 2009). This is significant because membrane-bound mucus may exist to protect extremely sensitive structures, such as olfactory sensory neurons (OSNs), that would be destroyed altogether if the entire mucus layer were lost.

While there is substantial understanding of mucin expression in the sinonasal respiratory epithelium, there are few studies of their expression in the olfactory epithelium that makes up 3% of human and 50% of mouse nasal mucosal surface area (Harkema et al. 2011; Solbu and Holen 2012). Within this epithelium, the cilia of olfactory sensory neurons contain receptors that detect odorant molecules within the overlying mucus layer (Buck 2005; Axel 2005). Odorant binding to these receptors initiates a second messenger signal transduction process that results in generation of action potentials (Restrepo et al. 1996; Gold 1999) traveling to second order neurons in the olfactory bulb. We hypothesize that expression of mucins in the mucus overlying the olfactory epithelium differs from their expression in the respiratory epithelium because of the inherently different functions of these epithelia. For instance, the abundant expression of mucins in the OE mucus may supplement the role that odorant binding proteins, also found in OE mucus, play in odor transport and chemical modification of chemosensory stimuli (Heydel et al. 2013; Block et al. 2015). In addition, OSNs by virtue of their anatomic location, are a vulnerable target for external environmental insults, such as exposure to toxins or infectious agents (Dando et al. 2014; van Riel et al. 2015). Since mucins help protect against infection (Rose and Voynow 2006; Hattrup and Gendler 2008), mucin expression in the olfactory epithelium may differ from expression in the respiratory epithelium.

Though radiation therapy has well-documented effects on mucous membranes and clinically apparent effects on mucus character, we do not know if radiation alters the composition of mucin production in a similar manner to other inflammatory disease states (eg, chronic rhinosinusitis).

In this study, we address the primary question whether mucin expression differs between the respiratory and the olfactory epithelium and as a secondary objective, how radiation alters mucin expression.

## Materials and Methods

### Animals

Nasal epithelia were obtained from two to four month old wild type C57/BL6 or OMP-ChR2-YFP C57BL/6 mice (Li et al. 2014). Mice were kept in the National Institutes of Health-approved animal facility of the University of Colorado Anschutz Medical Center. They were given food and water *ad libitum*, and were maintained in a 12 h light/dark cycle. All procedures were in compliance with the University of Colorado Anschutz Medical Campus Institutional Animal Care and Use Committee (IACUC).

### Tissue Preparation

Mice were anesthetized with fresh Avertin and perfused transcardially with 4% paraformaldehyde in 0.1 M phosphate buffer (PFA) as described by Nguyen and colleagues (Nguyen et al. 2012). All nasal tissue, including septum, turbinates and olfactory bulbs were resected en bloc and placed in 4% PFA. Tissue was cryoprotected with 20% sucrose in 0.1 phosphate buffer solution. Pairs of irradiated and control tissue were then transferred to cutting block molds, embedded in OCT and frozen to −20°C. We placed control and irradiated mouse tissue in the same cutting block to ensure consistency between the two groups during the IHC staining component and image acquisition process. Coronal sections of 12 μm were obtained in an anterior to posterior fashion using a microtome (Leica Biosystems, Buffalo Grove, IL) and thaw-mounted to glass slides, which were then stored at −20°C until staining.

### Immunohistochemistry

We followed a previously described protocol (Lopez et al. 2014) where samples were allowed to equilibrate to room temperature then washed three times in 0.1 M phosphate buffer solution (PBS). They were then exposed to a blocking solution (0.2 M phosphate buffer (PB), 0.05 M NaCl, 0.3% triton X-100, 3% BSA, 3% normal donkey serum (NDS)) for two hours at room temperature. Primary antibodies diluted in blocking solution were allowed to incubate for two days at 4°C. All primary antibodies were diluted to 1:300 and consisted of the following: rabbit anti-MUC1 (ab15481, Abcam); mouse anti-MUC2 (Ccp58: sc-7314 Santa Cruz Biotechnology); mouse anti-MUC5AC (MA5-12178, Thermo); rabbit anti-MUC5B (H-300: sc-20119 Santa Cruz Biotechnology); and goat anti-CNGA2 (SC-13700, Santa Cruz Biotechnology). Anti-CNGA2 was applied to all slides to differentiate olfactory epithelium (OE) from respiratory epithelium (RE), while only one anti-mucin was applied per slide. After incubation with the primary antibodies, slides were then washed in 0.1M PBS for 30 minutes with 3 changes of solution. Secondary antibodies were diluted in blocking buffer (1:1000) and incubated at room temperature for 2 hours then washed in PBS as above. Secondary antibodies consisted of anti-rabbit (A-21206 Alexa 488, A-10042 Alexa 568), anti-mouse (A-21202 Alexa 488, A-1137 Alexa 568), and anti-goat (A-11055 Alexa 488, A-1157 Alexa 568). Control slides were stained with secondary antibody, but no primary antibody, and no labeling was seen. Samples were counterstained with 1:5000 DRAQ5 (5mM) (ab108410, Abcam) for 10 minutes, washed as above, and mounted with Fluoromount G and coverslips.

### Image Acquisition and Processing

Samples were viewed and images acquired with a Leica TCS SP5 laser scanning confocal microscope with 10X and 20X air objectives and a 63X oil immersion objective (NA’s 0.4, 0.7, and 1.4, respectively). Leica Application Suite (LAS) AF was used to acquire z-stack images consisting of 12 to 20 1μm thick planes and saved as LIF files. ImageJ (Rasband 1994) with the Fiji (Schindelin et al. 2012) graphical user interface was used to generate maximum intensity projections from the confocal z-stacks. Staining patterns of each mucin with respect to OE and RE were visually observed in a blinded fashion for control and irradiated mouse tissue. Only unenhanced images from slides that showed both OE and RE within the same sample were used for quantitative analysis, which was performed by creating linear regions of interest (ROI) that isolated OE or RE as determined by CNGA2 staining. The plot profile analysis tool was used to extract single pixel fluorescence intensity values ranging from 0 to 255 along the lengths of each ROI. A two-tailed paired t-test was employed when comparing immunofluorescence intensity values between OE and RE from the same animal. To generate axial maximum intensity projections (Figure 3), the confocal datasets were loaded into MATLAB using our in-house toolbox, *imstack* (The MathWorks, Natick, MA). The datasets were then rotated around the axial axis using the *permute* function. Individual axial sections were selected for display using the *imstack* toolbox. Pixel dimensions were scaled along the z-axis using the *imresize* function.

### Antibody validation (Supplementary Figure 1)

#### Cell culture and immunocytochemistry

TSA-201 cells cultured at ~60% confluency were transfected with 1 ug/mL human MUC1 in pCMV3-GFPSpark (Sino Biological, Collegeville, PA; MG50877ACG) using Lipofectamine 3000 (Thermo Fisher). 48 hours post transfection, cells were washed with PBS and fixed with 4% PFA for 30 minutes at room temperature. After fixation, cells were washed with PBS and incubated in blocking buffer (0.3% Triton-X100, 1% BSA, and 2% normal donkey serum in 0.1M phosphate buffer (29 mM NaH2PO4, 75 mM Na2HPO4, pH 7.2-7.4)) for 1 hour at room temperature. MUC1 primary antisera (Abcam, ab15481, 1:300) was diluted in blocking buffer and applied overnight at 4C. Secondary antisera were applied (Alexa-568 donkey anti-rabbit, Invitrogen A10042, 1:800) for 2 hours at room temperature. Negative controls consisted of untransfected cells processed as above and transfected cells processed with omission of the primary antisera. After washing, cells were counterstained with DAPI (0.5 ug/mL) and imaged using Ocular software (QImaging) controlling a CCD camera (QImaging Retiga R3) connected to an Olympus IX71 inverted microscope with 40× air objective (NA 0.6). Negative control images were collected using the same acquisition settings.

**Figure 1.**
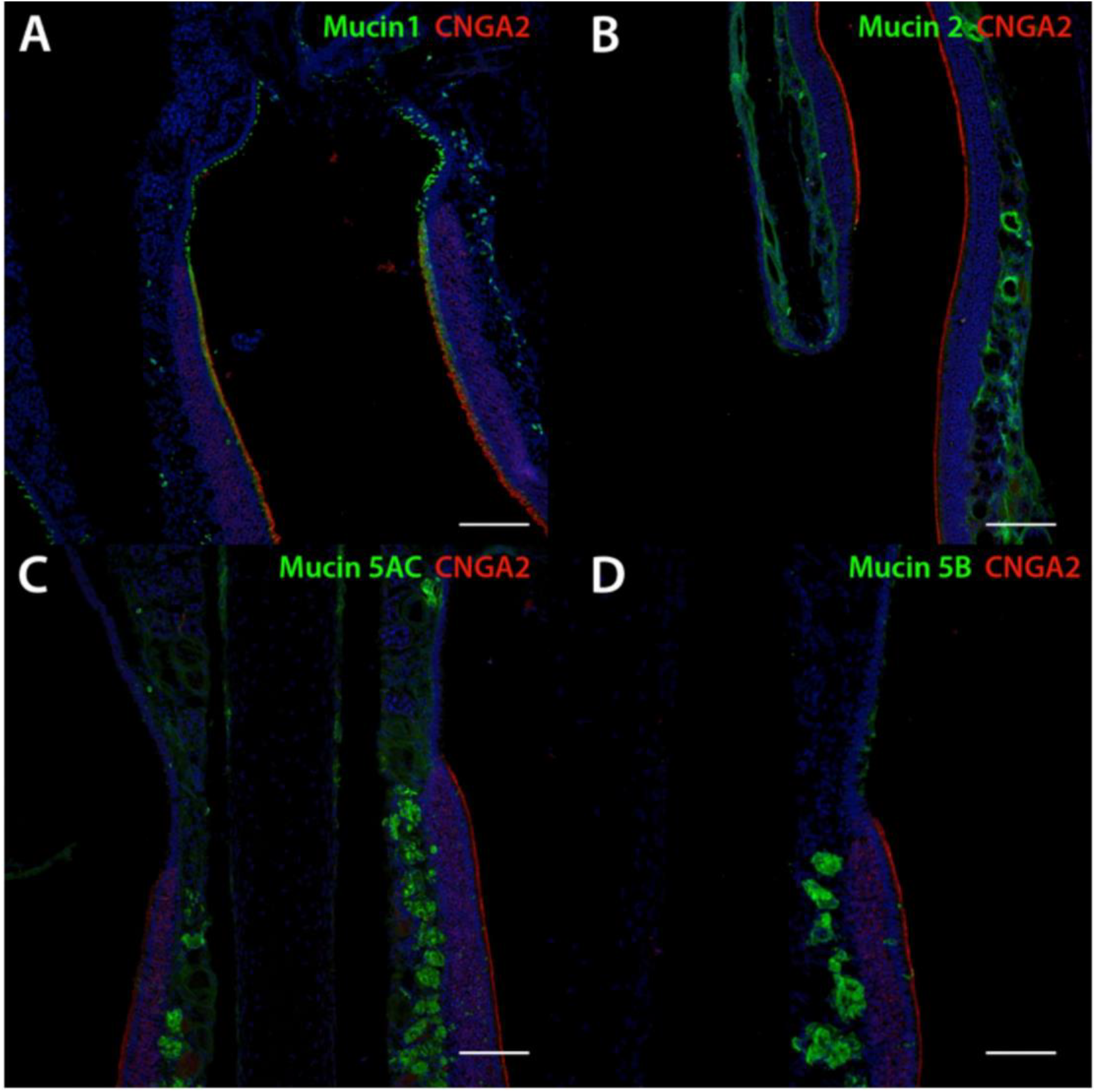
Transitions between olfactory epithelium (OE) and respiratory epithelium (RE) in mouse nasal tissue. CNGA2 (red), a marker specific to neurocilia of olfactory sensory neurons indicates portions of olfactory epithelium. A: Mucin 1 (green) is located uniformly across a continuous layer of sustentacular cells within the OE whereas in the RE scattered individual surface epithelial cells express it. B: Mucin 2 (green) is expressed uniformly within lamina propria connective tissue below both OE and RE. C: Mucin 5AC is expressed by submucosal glands and to a lesser degree within lamina propria connective tissue below both OE and RE. The density of positively staining glands was much higher in the OE. D: Mucin 5B was expressed intensely by submucosal Bowman’s glands within the OE, but within the RE goblet cells expressed mucin 5B. *Scale bars are 100 microns in all images*.

#### Image analysis

Raw images were imported to ImageJ (NIH, Bethesda, MD) where regions of interest were drawn around cells as identified with DAPI and/or a brightfield image. Background-subtracted fluorescence intensity for both GFP and Alexa-568 signals was measured and visualized on a scatterplot using SigmaPlot (Systat Studios).

#### Western Blot

Adult C57 mice were euthanized with CO2 and pooled respiratory and olfactory epithelial tissues were collected in T-PER (Thermo Fisher) protein extraction reagent. Tissue was gently homogenized using a hand-held blender before centrifugation at 500g for 10 minutes to remove nuclei and cell debris. Protein concentration was calculated using Pierce BCA protein assay kit (Thermo Fisher). 30 µg of protein was loaded on a 10% SDS-PAGE gel and separated by electrophoresis. Proteins were transferred to a nitrocellulose membrane and blocked with 5% non-fat milk in TBST. Primary antisera (MUC2, sc-73154; MUC5AC, MA5-12178; MUC5B, sc-20119) were diluted 1:1000 in TBST and applied overnight to separate blots at 4C. After thorough washing with TBST, HRP-conjugated secondary antibodies were applied for 1 hour at room temperature. HRP was detected using Pierce ECL substrate (Thermo Fisher) and imaged with a ChemiDoc XRS (Biorad). No-primary controls were run in parallel and exposed twice as long to ensure no non-specific signals. Blot images were acquired individually.

## Results

### Mucins are differentially expressed in mouse olfactory and respiratory epithelia

OE was identified and distinguished from RE by the presence of CNGA2, an ion channel subunit unique to OE (Liman and Buck 1994). Of the four mucins we studied, only mucin *2* showed a similar staining pattern in both OE and RE tissue (Figure 1). Mucin 2 is a secreted mucin, and we found it clustered in mucus overlying the luminal areas of samples where the secreted mucus layer survived the sample preparation process. We also observed mucin 2 diffusely expressed in the connective tissue of the lamina propria underlying both OE and RE. Quantitatively, there was a significant difference in the immunofluorescence intensity of mucin 2 between the OE and RE lamina propria (p < 0.01; Figure 2); however this difference was small and may not be functionally significant. Unlike the other secreted mucins (mucins 5AC and 5B), mucin 2 was not seen in glandular structures.

**Figure 2.**
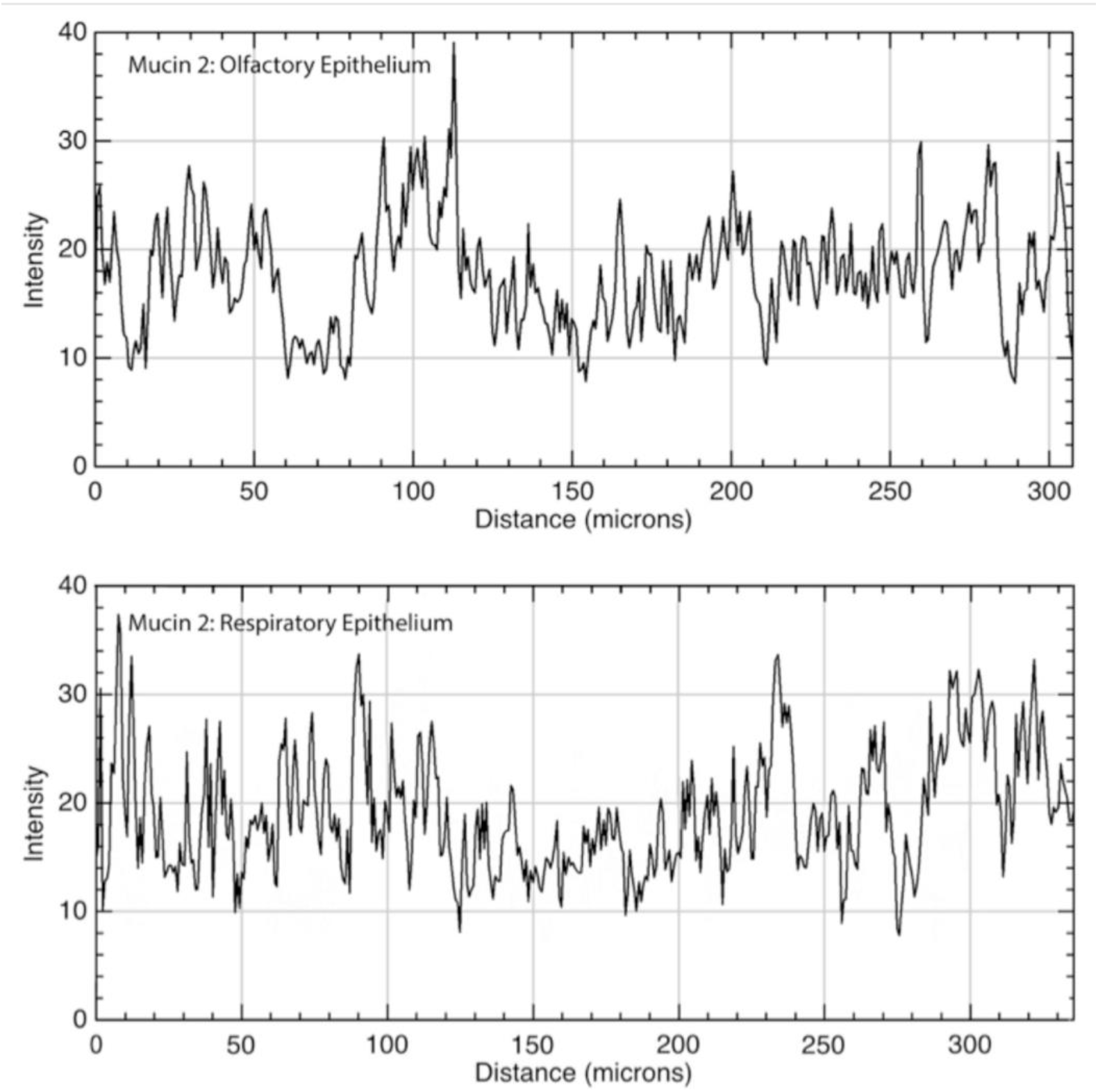
Immunofluorescence quantitation of mucin 2 in the lamina propria underlying olfactory and respiratory epitheliia. Mucin 2 was diffusely present within the lamina propria of both the olfactory (top panel) and respiratory (bottom panel) epithelia. X-axis is the distance moving along a linear region of interest within the lamina propria, parallel to the epithelial layer; Y-axis is intensity of immunofluorescence with possible values ranging from of 0-255. Mean OE versus RE immunofluorescence was 19.0 vs 17.7, respectively (p<0.01).

We observed differential expression of mucins 1, 5AC and 5B between OE and RE. *Mucin 1* is membrane-bound and was observed in a lattice-like pattern at the level of the OSN dendrites (Figure 3). The “spaces” in the lattice where mucin 1 is absent were measured at 1.5 (SD 0.21) microns in diameter, which corresponds to the size of dendrites or dendritic knobs from which olfactory cilia project (Kwon et al. 2009; Oberland et al. 2015; Ma et al. 2003). Within the OE, mucin 1 shows a consistent, linear, 1.1 (SD 0.11) micron thick layer between the cilia and their cell bodies. In contrast, within the RE, mucin 1 demonstrates a patchy pattern of expression that upon higher magnification was limited to scattered individual cells. Immunofluorescence intensity plots demonstrated a relatively consistent value across the OE (mean 36, SD 13) whereas the RE showed high intensity peaks that coincided with individual cells separated by troughs of minimal immunofluorescence (Figure 4).

**Figure 3.**
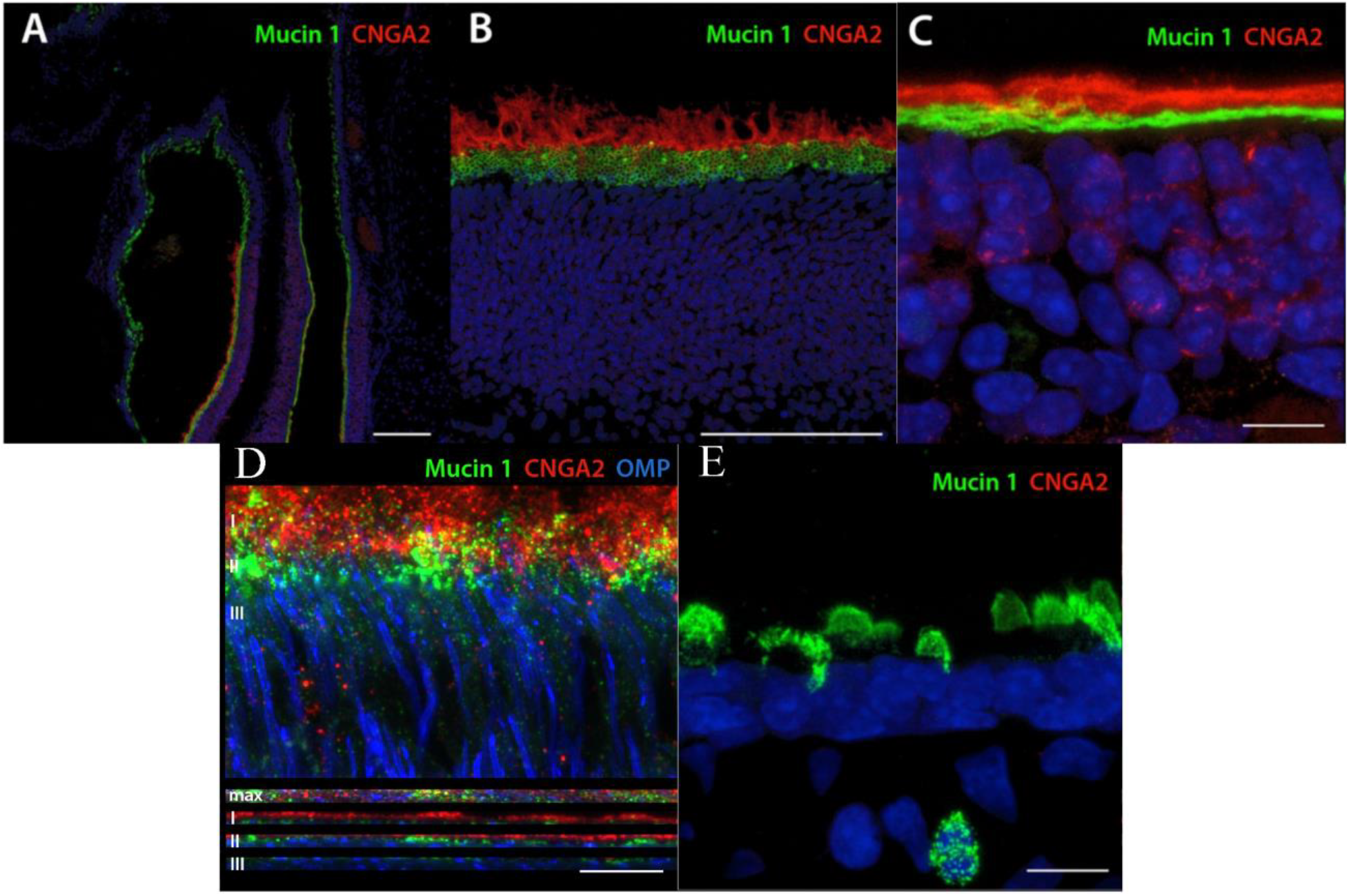
Mucin 1 differential staining between olfactory epithelium and respiratory epithelium. A: Mucin 1 is expressed in a uniform layer over the OE whereas in the RE it is expressed by scattered individual cells. B and C: In the OE, mucin 1 lies at the base of the neurocilia. When viewed in oblique section (B), mucin 1 exhibits a lattice-like staining pattern suggestive of perforations by OSN dendritic knobs, whereas in true orthogonal section (C), a complete layer is seen at the base suggesting it may be produced and secreted into this layer by the sustentacular cells. D: OMP-ChR2-YFP knock-in mice were used to examine the relationship between OSNs (blue), Mucin 1 (green), and CNGA2 (red). Mucin 1 is found in a layer just above the sustentacular cells, and beneath the cilia, as olfactory dendrites coursing through the OE are seen. Coronal projections at varying depths (I,II, and III) reveals olfactory dendrites penetrating into the mucin layer beneath the neurocilia. E: In RE, MUC1 is expressed on the surfaces of scattered secretory cells and extends into the luminal space. *Scale bars are 100 microns in A & B; 10 microns in C, D & E*.

**Figure 4.**
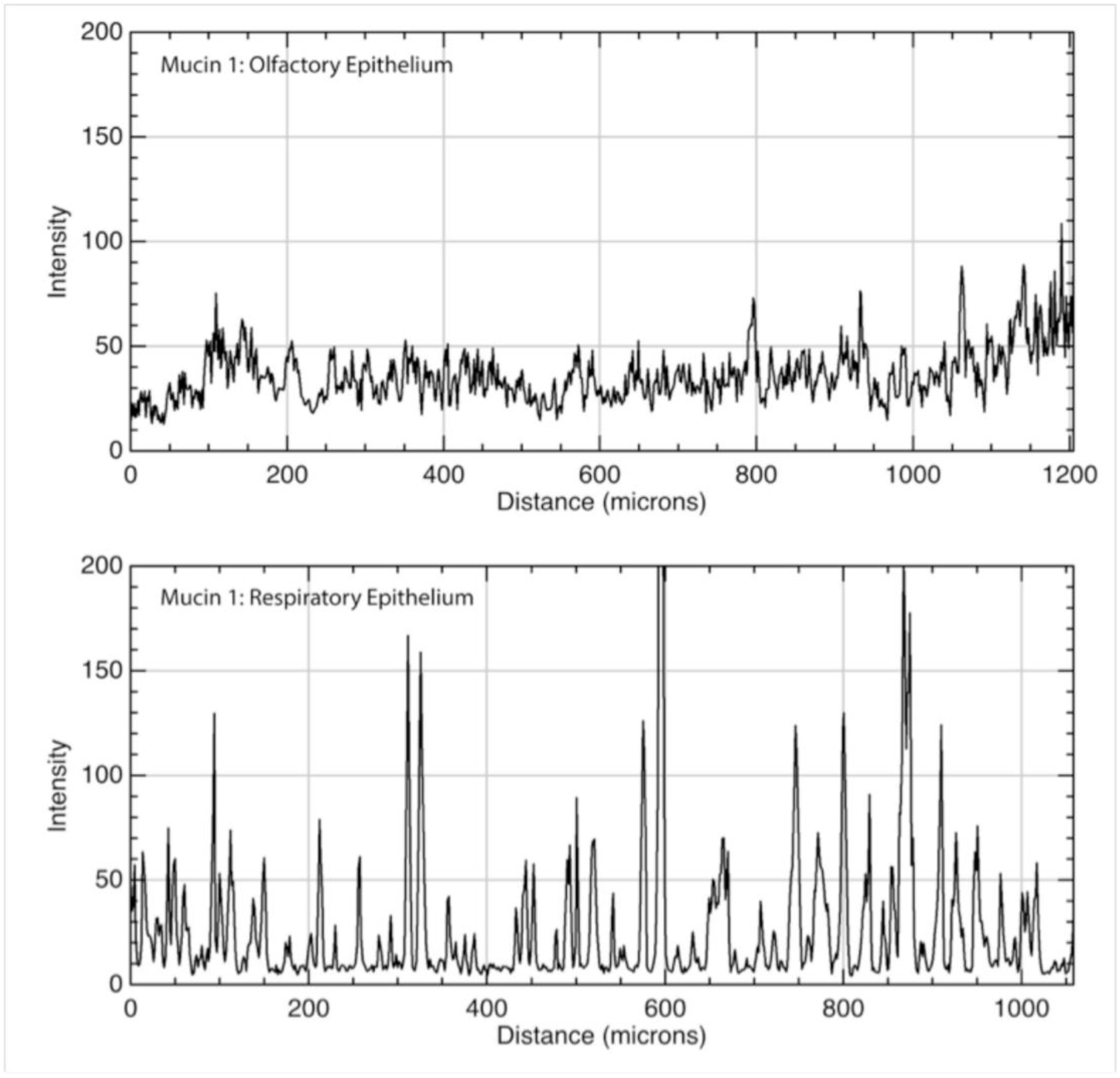
Immunofluorescence quantitation of mucin 1 in olfactory and respiratory epithelium. **Top**: Mucin 1 was diffusely present along the apical surface of olfactory epithelial cells. **Bottom**: In the respiratory epithelium, it was present on the apical surfaces of individual goblet cells, resulting in a series of immunofluorescence peaks. X-axis is distance along a linear region of interest consisting of the apical surface of the epithelium; Y-axis is intensity of immunofluorescence with possible values ranging from of 0-255. Values indicate consistent moderate expression of the membrane-bound protein throughout the OE, versus intermittent areas of high expression in RE consistent with goblet-cell co-localization.

Mucin 5AC is found in the mucus layer overlying OSN cilia and portions of the RE where it survived the sample preparation process, consistent with its known action as a secreted mucin. Connective tissue within the lamina propria stained diffusely positive for mucin 5AC at a low level (mean 5.2, SD 3.1). Subepithelial glands with ducts from the lamina propria to the epithelial surface stained positive for mucin 5AC at a higher level than the surrounding connective tissue (mean 35.5, SD 21.5), and the frequency of these glands was substantially greater in the OE versus RE of our samples (Figures 1 and 5). This is consistent with previous research demonstrating that Bowman’s glands secrete mucin 5AC in rodents (Solbu and Holen 2012).

**Figure 5.**
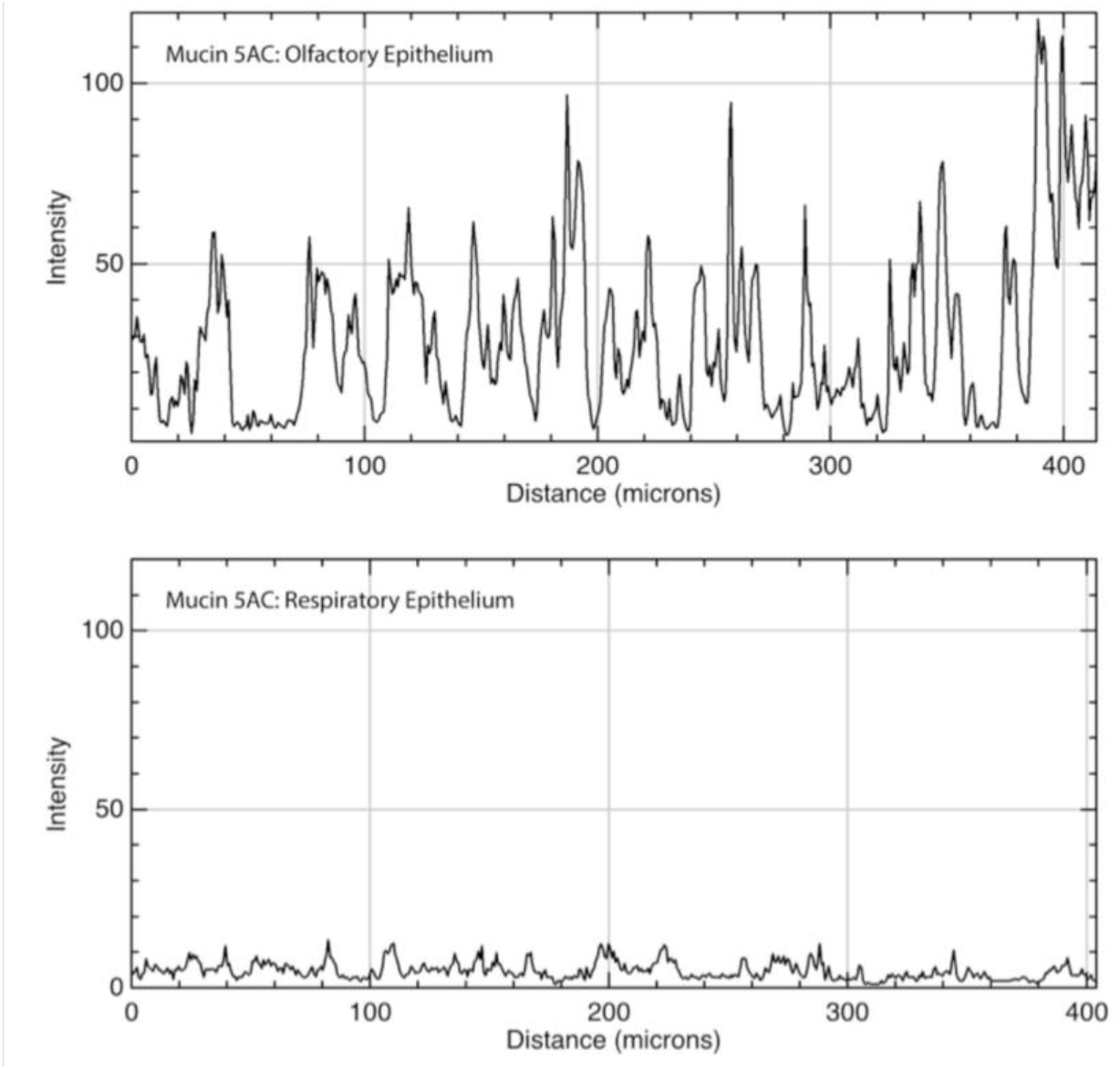
Immunofluorescence quantitation of mucin 5AC in olfactory and respiratory epithelium. **Top**: Mucin 5AC was present primarily in submucosal glands within the olfactory epithelium and to a much lesser degree within the surrounding lamina propria connective tissue. **Bottom**: In the respiratory epithelium, there was a low level of immunofluorescence within the connective tissue and very few glands expressing the protein (none are present in the sample above). X-axis is distance along a linear region of interest consisting of the submucosa, parallel to the epithelium; Y-axis is intensity of immunofluorescence with possible values ranging from of 0-255.

Mucin 5B is also a secreted protein and was present within clusters of mucus on the luminal surface of both OE and RE. In contrast to mucin 5AC, mucin 5B did not appear within lamina propria connective tissue and also had a different pattern of glandular staining. Within the OE, mucin 5B-producing glands existed submucosally with ducts opening to the epithelial surface, and these glands appeared to have an acinar structure with a central lumen surrounded by a ring of secretory cells (image not shown). In contrast, epithelial rather than submucosal cells expressed mucin 5B in the RE. These single cells were spaced apart, usually within 15 microns of each other, and at low power magnification had a granular staining pattern within the epithelium (Figures 6 and 7). Because mucin 5B expression was subepithelial in the OE and superficial in the RE, this pattern easily differentiated OE from RE.

**Figure 6.**
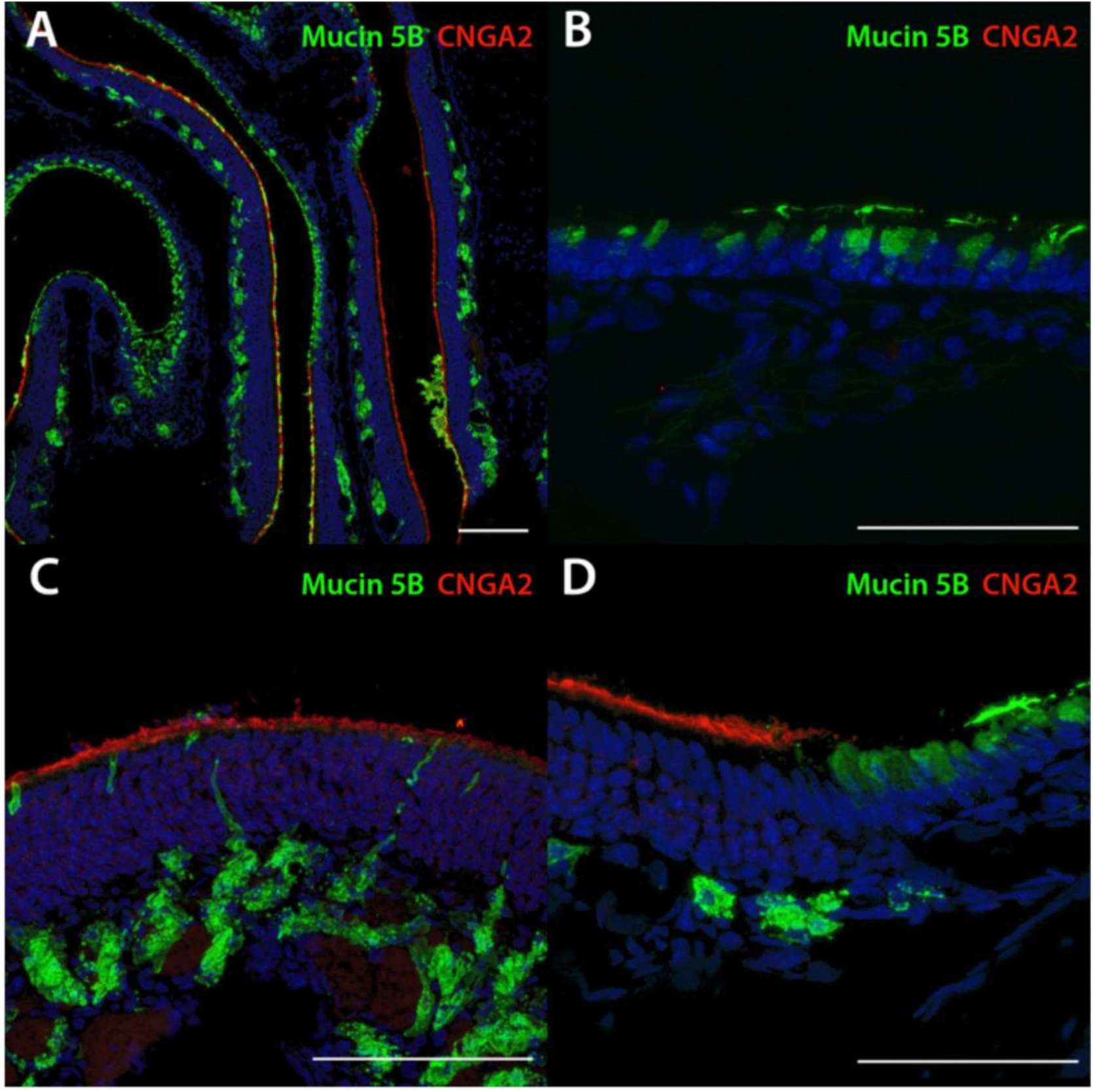
Mucin 5B differential staining between olfactory epithelium and respiratory epithelium. A: In the OE, mucin 5B is located in submucosal glands while in the RE it is secreted by surface goblet cells. B: Magnification of RE showing goblet cells secreting directly to the epithelial surface. Note the vesicles filled with mucin oriented to the apical side while the nuclei are located basolaterally. C: Magnification of OE showing submucosal glands with ducts to the luminal surface. D: Transition between OE (left) and RE (right). *Scale bars are 100 microns in all images*.

**Figure 7.**
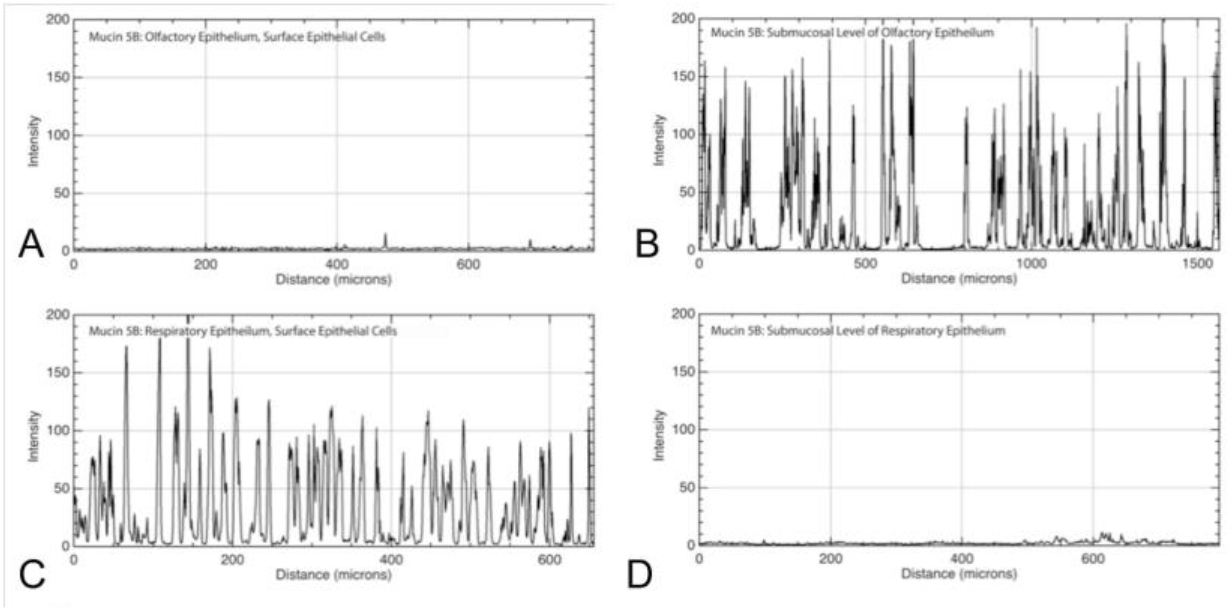
Immunofluorescence quantitation of mucin 5B at the epithelial and submucosal level of olfactory and respiratory epithelia. Mucin 5B exhibited almost no immunofluorescence in the epithelial layer (Panel A), but was present primarily in submucosal glands of the olfactory epithelium as evidenced by the strong peaks of immunofluorescence (Panel B). Individual goblet cells throughout the surface of the respiratory epithelium expressed high levels of mucin 5B, resulting in immunofluorescence peaks (Panel C). There was virtually no mucin 5B in the submucosal level of respiratory epithelium (Panel D). X-axis is distance along a linear region of interest; Y-axis is intensity of immunofluorescence with possible values ranging from of 0-255.

### Radiation did not affect mucin expression at one week

We analyzed mucin expression one week after mice received 8 gy of radiation to the anterior cranium in a single dose, in the same fashion. Comparing control to irradiated tissues using similar qualitative and quantitative testing (unpaired t-test) as above, we did not find any consistent changes in mucin expression between the irradiated and non-irradiated samples (data not shown).

## Discussion

### Comparison with human nasal mucosa

There is evidence that murine nasal mucosa approximates that of humans (Harkema et al. 2011), making it a good model for study of mucins in the olfactory epithelium. The gel-forming secreted mucins 2, 5AC and 5B we studied as well as mucin 6 are encoded by the same gene cluster located on chromosome 11p15.5 in humans and chromosome 7 band F5 in mice, and each of these mucins has been shown to be highly conserved across species (Escande et al. 2004; Pigny et al. 1996; Desseyn and Laine 2003; Chen et al. 2001; Linden et al. 2008). Patterns of expression within respiratory epithelium of the larynx, trachea, and lungs also appear similar between mice and humans though to date, no study has analyzed patterns of mucin isoform expression in mouse nasal mucosa. Multiple studies have analyzed nasal mucin isoform expression in humans and have found that mucin 5AC is weakly expressed by surface goblet cells of the nasal epithelium in healthy samples while mucin 5B is produced by both goblet cells and submucosal glands (Groneberg et al. 2003; Ali et al. 2005; Ali et al. 2002; Ding and Zheng 2007; Kim et al. 2004; Fahy and Dickey 2010). In our mouse study, we found mucin 5AC ubiquitously in submucosal connective tissue as well as submucosal glands that were clustered more closely in OE than RE. We did not observe mucin 5AC production within the epithelium in structures consistent with goblet cells. In contrast, mucin 5B was produced by cells morphologically resembling goblet cells and submucosal glands consistent with human studies with one important caveat. We observed a differential expression of mucin 5B based on the overlying mucosal tissue type. In RE, mucin 5B was confined to goblet cells while in OE only submucosal glands produced mucin 5B. The acinar structure of the glands expressing mucin 5B is consistent with Bowman’s glands (Nomura et al. 2004; Solbu and Holen 2012). The difference in surface versus submucosal expression was consistent and could be used to differentiate these two tissue types. Airway secretion of mucin 5B has been shown to be essential to murine life (Fahy and Dickey 2010), and this may explain why mucin 5B was produced across both epithelial types. Due to the density of neurons competing for access to the luminal surface of OE, mucin 5B production may be forced to go “underground” to the submucosa where there is room enough to accommodate glands. Without olfactory neurons occupying the epithelial surface in the RE, there is room for production of this essential airway mucin within the luminal cells.

### Mucin 1 expression is located to a dendritic layer under the cilia of OSNs

We found that mucin 1 appears to tightly surround the dendrites or dendritic knobs of olfactory neurons (Figure 3). Mucin 1 has been postulated to defend against pathogens and toxins via a range of mechanisms, which is significant given access to the intracranial compartment presented by olfactory neurons (Dando et al. 2014; Nguyen et al. 2011; McAuley et al. 2007; Guang et al. 2010). Mucin 1 contains a large protein core with dense sugar chains that interfere with pathogen and toxin binding by denying them access to receptors on underlying cells, blocking them from penetrating inter-cellular spaces, and disrupting pathogen adhesion via a strong negative charge (Hattrup and Gendler 2008), Additionally, mucin 1 mimics some cellular pathogen binding sites, and though mucin 1 is tethered to the cell membrane, it has the ability to slough off and shed the bound pathogens (Linden et al. 2008; Guang et al. 2010; McGuckin et al. 2007; Linden et al. 2009). The intracellular portion of mucin 1 affects cellular survival in response to pathogens by down-regulating the immune system to avoid excessive inflammation that would degrade the integrity of the mucosal barrier (Hattrup and Gendler 2008; Groneberg et al. 2003; McAuley et al. 2011; Kim and Lillehoj 2008; Kyo et al. 2012; Ueno et al. 2008). Mucin 1 has also been shown to counteract genotoxins that trigger apoptosis, which is useful in responding to some bacterial attacks, but may be detrimental in others. For instance, mucin 1 is considered an oncoprotein, in part due to its ability to promote cancer cell survival and resistance to chemotherapies by counteracting signals that would trigger apoptosis in healthy cells (Linden et al. 2008; McAuley et al. 2011; Kufe 2009).

In our olfactory epithelial samples, the OSN cilia projected above the mucin 1 layer by 2.4 micrometers (SD 0.3 micrometers). Although we did not co-stain for different mucins, comparison between images indicates that the mucin 1 layer in the OE is beneath the layer of mucus containing mucins 5AC and 5B resting on top of the olfactory cilia. This supports the model of a slippage plane formed by distinct membrane bound and secreted mucin layers. This is significant because membrane-bound mucus may exist to protect extremely sensitive structures, such as olfactory sensory neurons, that could be susceptible to damage if the entire mucus layer were lost. Mucins give the mucus layer its unique physical properties of thixotropy which allows mucus to slide smoothly when exposed to high shear stress like coughing or sneezing but then become less mobile and gel-like under low stress (Lai et al. 2009). This may be another method of conferring protection to the underlying sensory neurons by diffusing high velocity stress within the mucus layer rather than transferring it to the underlying epithelium, such as occurs in blunt head trauma.

Our findings of mucin 1 expression in the OE, and the associated slippage plane of overlying mucins, suggest that these mucins may protect the delicate OSNs, and indicates a rationale for continued research.

### Effects of radiation

We did not observe any effects of radiation exposure on mucin expression at the one-week time point. We selected a single 8gy dose because this has been shown to alter cellular function within mouse olfactory and taste mucosa without resulting in excess mortality (Nguyen et al. 2012; Cunha et al. 2012). At one-week post irradiation, Cunha et al. reported changes in mouse nasal mucosal morphology as well as olfactory performance as measured by behavioral testing. Specifically, they found that 8 Gy induced a decrease in OSN proliferation but an increase in sustentacular cell proliferation. CD15, a marker for Bowman’s glands, appeared between OSNs after irradiation. We observed that in portions of one irradiated specimen, mucin 2 was highly expressed within the OE epithelial layer in a strand-like fashion suggesting it interdigitated between cells while in the other irradiated and control mice mucin 2 was confined to the lamina propria (figure not shown). This may parallel Cunha’s finding of increased CD15 between OSNs after radiation; however, our observation was not sufficiently consistent to form any conclusions. Cunha also reported a disruption in the basal lamina at the 5-week time point compared to 24 hours post irradiation, though no intermediate time points were reported. We did not observe any changes in the basal lamina at one week. Given that other studies have shown alterations in mucin production in response to sinonasal irritation (Ali and Pearson 2007; Fahy and Dickey 2010), it is possible that one-week post-treatment is too soon to see any detrimental effects on mucin production. Based on our results, it appears that 1-week was either too soon for radiation to effect mucin expression or that mucins may not substantially affected by radiation.

### Limitations & directions for further research

We designed our radiation model with the intent that it would help us better understand the pathophysiology that underlies radiation damage to mucosal tissues observed in humans. Our single dose model was similar to that employed in other animal studies, but head and neck cancer patients undergo fractionated radiation rather than single exposures. Currently available literature does not shed any light on the comparison of single dose versus fractionated radiation effects on airway mucosa. We used a sample size of six mice in the radiation group, and results from control mice were consistent across samples and similar to data from healthy human tissue studies, a reassuring finding. Only one irradiated sample demonstrated an inflammatory response with mucin 2 expressed within the epithelial layer. It is possible that the experiment was underpowered, or alternatively, that delayed post-treatment time points may show differences.

This study demonstrated differential mucin expression between the olfactory and respiratory epithelia, and mucin 1 in particular seems to have a unique distribution suggestive of a special role within the olfactory epithelium. Further research will test mucin 1’s protective effects on susceptibility to pathogen infection (Nguyen et al. 2011), and its ability to protect OSNs from physical injury.

## Conclusion

Olfactory epithelium is unique from respiratory epithelium with regard to expression of mucins 1, 5AC and 5B. The physical relationship of mucin 1 with olfactory sensory neurons and its known role in mucosal defense suggests it may serve to protect OSNs from injury, and by extension, the central nervous system from environmental insult. Olfactory epithelium expresses more mucin 5AC than does respiratory epithelium, whereas mucin 5B is ubiquitous across both epithelial types, and this pattern consistently distinguishes OE from RE. Lastly, radiation did not appear to affect mucin expression at the one-week time point.

## Level of Evidence

N/A (Basic Science Research)

## Abbreviations (do not use any in title or abstract)

CNG: cyclic nucleotide-gated
IHC: immunohistochemical
NDS: normal donkey serum
OE: olfactory epithelium
OSN: olfactory sensory neuron
PB: phosphate buffer
PBS: phosphate buffer solution
PFA: paraformaldehyde in 0.1 M phosphate buffer
RE: respiratory epithelium

## Acknowledgements

We would like to thank Tom Finger, PhD, for valuable advice in fluorescence microscopy and imaging, and review of early versions of this manuscript.

---

**Supplementary Figure 1.**
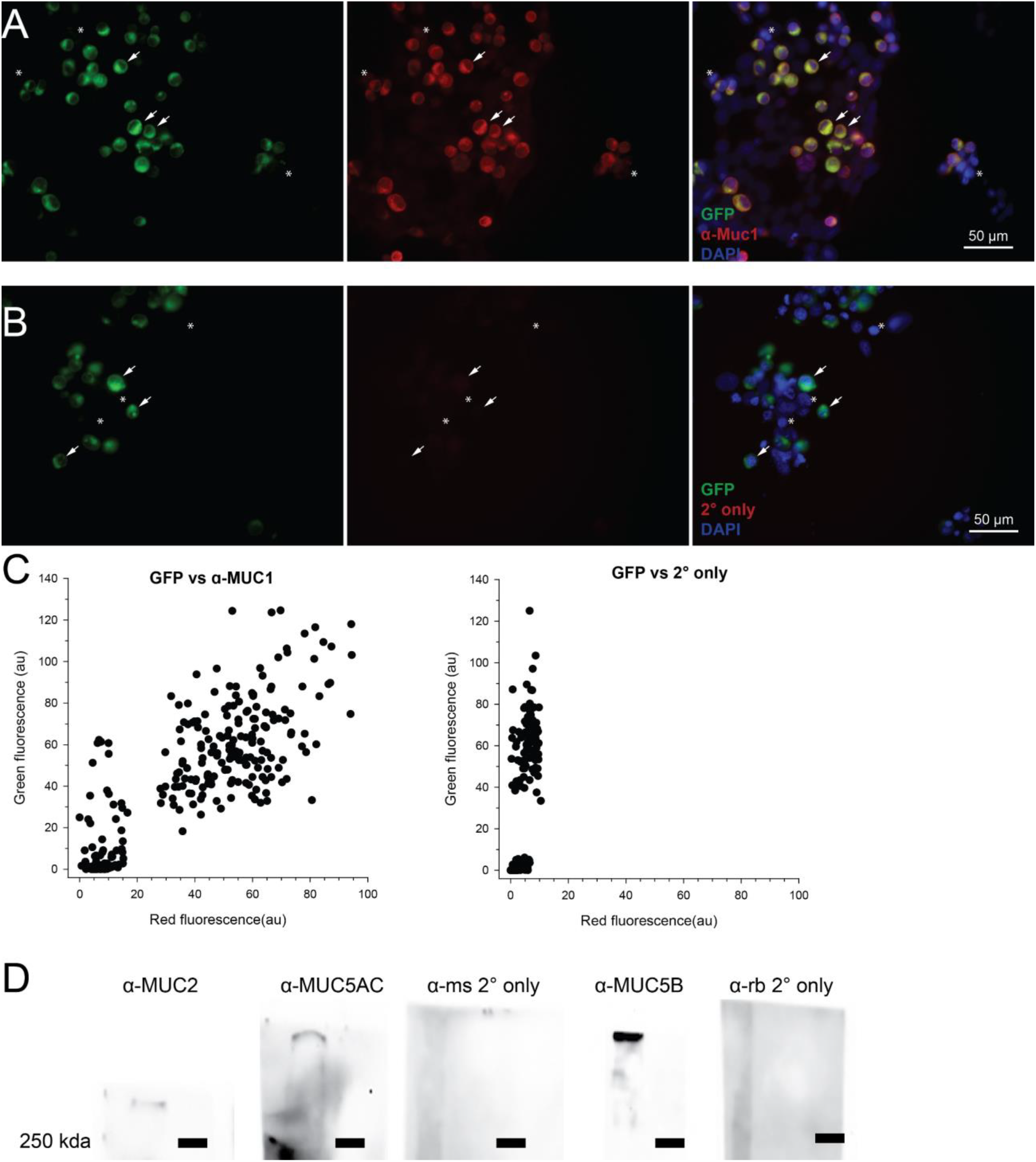
Validation of MUC1 antibodies. A-B. TSA-201 cells transfected with human MUC1 in pCMV-GFPSpark. Transfection was confirmed by presence of GFPSpark. Cells were incubated with (A) or without (B) primary antibody to MUC1 (ab15481). Arrows show example transfected cells and asterisks show example non-transfected cells. Transfected cells showed a strong immunoreactivity to MUC1 while non-transfected cells did not. Image acquisition settings between panels A and B were kept constant. C. Scatter plot showing average pixel intensity of green and red channels. Each dot represents a region of interest drawn around a cell. With very few exceptions, transfected cells were highly immunoreactive for MUC1 while non-transfected cells were not. D. Western blots using MUC2, MUC5AC and MUC5B antisera on mouse nasal epithelial homogenates showed immunoreactive bands >250 kDA. Mucins are large glycoproteins with predicted molecular weights between 300 and 600 kDA. Omission of primary antisera showed no signal. Black bars denote 250 kDA ladder. Blot images were acquired individually.

